# How strongly does appetite counter weight loss? Quantification of the homeostatic control of human energy intake

**DOI:** 10.1101/051045

**Authors:** David Polidori, Arjun Sanghvi, Randy Seeley, Kevin D. Hall

**Author notes:** To whom correspondence should be addressed: Kevin D. Hall, PhD, National Institute of Diabetes & Digestive & Kidney Diseases, National Institutes of Health, 12A South Drive, Room 4007, Bethesda, MD 20892-5621.

## Abstract

**Objective:** To quantify the homeostatic feedback control of energy intake in response to long-term covert manipulation of energy balance in free-living humans.

**Methods:** We used a validated mathematical method to calculate energy intake changes during a 52 week placebo-controlled trial in 153 patients treated with canagliflozin, a sodium glucose co-transporter inhibitor that increases urinary glucose excretion thereby resulting in weight loss without patients being directly aware of the energy deficit. We analyzed the relationship between the body weight time course and the calculated energy intake changes using principles from engineering control theory.

**Results:** We discovered that weight loss leads to a proportional homeostatic drive to increase energy intake above baseline by ~100 kcal/day per kg of lost weight – an amount more than 3-fold larger than the corresponding energy expenditure adaptations.

**Conclusions:** While energy expenditure adaptations are often thought to be the main reason for slowing of weight loss and subsequent regain, feedback control of energy intake plays an even larger role and helps explain why long-term maintenance of a reduced body weight is so difficult.

**Funding:** This research was supported by the Intramural Research Program of the NIH, National Institute of Diabetes & Digestive & Kidney Diseases, using data from a study sponsored by Janssen Research & Development, LLC.

**Disclosure:** D.P. is a full-time employee of Janssen Research & Development, LLC. K.D.H. reports patent pending on a method of personalized dynamic feedback control of body weight (US Patent Application No. 13/754,058; assigned to the NIH) and has received funding from the Nutrition Science Initiative to investigate the effects of ketogenic diets on human energy expenditure. R.S. is a paid consultant for Janssen, Novo Nordisk, Takeda, Daichii Sankyo, Novartis, Pfizer, Nestle, Circuit Therapeutics and Ethicon. R.S., also has received research support from Novo Nordisk, Ethicon, Sanofi and Boehringer Ingelheim. A.S. reports no conflicts of interest.

**What is already known about this subject?:**

- Human body weight is believed to be regulated by homeostatic feedback control of both energy intake and energy expenditure.
- Adaptations of energy expenditure to weight loss have been well-established, but the homeostatic control of energy intake has yet to be quantified.

**What this study adds:**

- We provide the first quantification of the homeostatic control of energy intake in free-living humans.
- The increase in energy intake per kg of weight lost is several-fold larger than the known energy expenditure adaptations.
- Homeostatic control of energy intake is likely the primary reason why it is difficult to achieve and sustain large weight losses.

## Introduction

Body weight is believed to be regulated by homeostatic control of both energy intake and energy expenditure. Several carefully controlled feeding experiments in humans have quantified how energy expenditure adapts in response to alterations of energy intake and body weight. For example, Leibel and colleagues found that energy expenditure changed by several hundreds of kcal/day thereby acting to resist weight changes (1). Such data have enabled the development of accurate mathematical models of energy expenditure and body weight dynamics in humans in response to given changes in energy intake (2). In contrast, energy intake adaptations have been difficult to accurately quantify in humans despite the widespread belief that homeostatic control of energy intake is critical for body weight regulation (3, 4, 5) and acts as part of a complex neurobiological system to determine overall human food intake behavior (6, 7).

Why has the assessment of human homeostatic energy intake control lagged the quantification of energy expenditure changes with weight loss? First, we lacked the ability to accurately measure changes in free-living energy intake in large numbers of people over extended time periods. While accurate energy intake measurements can be performed while subjects are housed in laboratory setting, such studies are typically of short duration and the artificial nature of the environment makes it difficult to translate the results to the real world (8, 9). Indeed, free-living energy intake is known to fluctuate widely from day to day and exhibits little short-term correlation with energy expenditure or body weight (10, 11). Therefore, observations over long time scales are required to investigate regulation of human energy balance and body weight (12) making laboratory based studies impractical.

Unfortunately, free-living subjects are notorious for being unable to provide accurate estimates of energy intake using self-report methods (13) and the expense and difficulty of employing objective biomarker methods severely limits their applicability (14). To address this important problem, we recently developed an inexpensive mathematical method for calculating energy intake changes in free-living subjects using repeated body weight data (15, 16). This method was validated using data from 140 free-living people who participated in a two year calorie restriction study where the mean calculated energy intake changes at all time points were within 40 kcal/day of measurements obtained using an expensive biomarker method (16).

The second impediment to quantifying energy intake control in humans is that we lacked an intervention that increases energy output without participants consciously knowing that this is occurring. Rather, most interventions that alter body weight or energy expenditure also evoke central responses that may mask the effect of weight changes per se on the homeostatic control of energy intake in humans. For example, engaging in an exercise program might increase energy expenditure and lead to weight loss, but exercise is a conscious behavior that doesn’t have an effective placebo control. Furthermore, exercise has a complex role in modulating appetite (17) and may induce compensatory changes in other components of total energy expenditure that are difficult to quantify. Therefore, changes in energy intake during an exercise program may not solely be due to homeostatic mechanisms and are likely to also involve conscious changes in behavior.

Here, we used data from a placebo-controlled trial in patients with type 2 diabetes who were treated for one year with canagliflozin, an inhibitor of sodium glucose transporter 2 (SGLT2), thereby increasing energy output in the form of urinary glucose excretion (UGE) (18). In patients with type 2 diabetes, treatment with canagliflozin at a dose of 300 mg/day increases mean daily UGE by approximately 90 g/day which is sustained at the same level throughout the duration of treatment (19). This leads to a net energy loss of ~360 kcal/day that occurs without directly altering central pathways controlling energy intake and without the patients being directly aware of the energy deficit. In other words, SGLT2 inhibition provides a novel way to perturb human energy balance that largely bypasses the volition of the subjects. Any observed increased energy intake countering the weight loss induced by SGLT2 inhibition therefore likely reflects the activity of the homeostatic feedback control system. We calculated the free-living energy intake changes in 153 patients treated with 300 mg/day canagliflozin over a 52 week trial using the measured body weight data and an assumed mean UGE of 90 g/day as inputs to a mathematical model that has recently been validated against an expensive biomarker method (16). We found that the homeostatic feedback control of energy intake in humans was proportional to the amount of weight lost and was substantially stronger than the control of energy expenditure. These findings have important implications for interpreting the results of obesity interventions.

## Methods

### Calculating changes in energy intake during canagliflozin treatment

We used measured body weight, BW, and baseline patient characteristics in the previously published placebo-controlled trail of canagliflozin (18) where compliance was monitored by pill counts. All of the subjects in the canagliflozin cohort who completed the study had compliance of at least 75% and all but 3 subjects in the placebo cohort who completed the study had >75% compliance. We calculated the changes in energy intake, Δ*EI*, for each subject using a validated mathematical method (16) using the following equation:

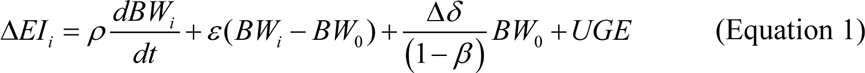

The inputs to the model were the change of body weight versus baseline over each interval, (*BW*_*i*_-*BW*_*0*_), and the moving average of the measured body weight time course was used to calculate the rate of change of body weight over each interval, *dBW*_*i*_/*dt*. The interval length was *t* = *(N-1)*\**T*, where *N* = 2 was the number of body weight measurements per interval and *T* = 52 was the number of days between measurements. Three subjects treated canagliflozin and 2 subjects treated with placebo had missing body weight data at one time point and linear interpolation was used to impute the missing data.

Equation 1 is a linearization of a mathematical model of adult body weight dynamics that was developed and validated using data obtained primarily from controlled feeding studies in adult humans with longitudinal measurements of changes in body composition as well as both resting and total energy expenditure (20, 21, 22). The model parameter *ρ* was the effective energy density associated with the body weight change:

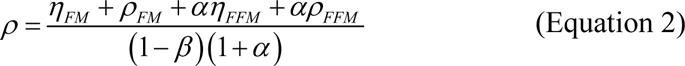

and *ɛ* defined how energy expenditure depends on body weight:

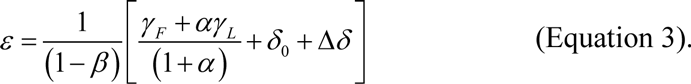

The model parameters *γ*_*FFM*_ and *γ*_*FM*_ are the regression coefficients relating resting metabolic rate to fat-free mass (FFM) and fat mass (FM), respectively. Parameters ρFM and ρFFM are the energy densities associated with changes in FM and FFM, respectively. Most physical activities involve locomotion and have an energy cost that is proportional to body weight for a given intensity and duration of activity (23). The baseline physical activity parameter was *δ*_*0*_ and Δ*δ* represents changes in physical activity that can be informed by objective measurements, if available. Without such measurements, physical activity changes are often assumed to be zero (Δ*δ*=0) with the realization that the calculated energy intake changes may be in error, especially at the individual level where substantial physical activity changes can occur throughout a study. Averaging over many individuals with the assumption that Δ*δ*=0 can also result in a biased mean energy intake change of the group, but our previous validation study demonstrated that this bias is likely to be <40 kcal/day when studying >100 individuals (16).

The parameter *β* accounts for the adaptation of energy expenditure during a diet perturbation, Δ*EI*, and was determined using data from eight human studies that measured changes in body composition as well as both total and resting energy expenditure before and after achieving a period of long-term stability at a lower weight (21). Parameters *η*_*FM*_ and *η*_*FFM*_ account for the biochemical cost of tissue deposition and turnover assuming that the change of FFM is primarily accounted for by body protein and its associated water (24).

The parameter *α* represents the relationship between changes of lean and fat mass: *α* = *dFFM/dFM* = *C/FM* where C = 10.4 kg is the Forbes parameter (25). This simple model of body composition change provides more accurate predictions than more complex models (26) and has been demonstrated to be consistent across different ethnic groups and sexes (27). For modest weight changes, *α* can be considered to be approximately constant with *FM* fixed at its initial value FM_0_. The larger the initial fat mass, FM_0_, the smaller the parameter *α*, as previously described (22). The parameter *UGE* represents the energy losses as a result of increased urinary excretion of glucose with canagliflozin treatment. Model parameter values are given in **Table 1**.

**Table 1.** Mathematical model parameters.

| Parameter | Value | Description |
| --- | --- | --- |
| $\gamma_{FM}$ | 3.2 kcal/kg/d | Energy expenditure rate of fat mass |
| $\gamma_{FFM}$ | 22 kcal/kg/d | Energy expenditure rate of fat free mass |
| $\delta_0$ | 10 kcal/kg/d | Physical activity at baseline |
| $\Delta\delta$ | 0 kcal/kg/d | Physical activity changes |
| $\eta_{FM}$ | 180 kcal/kg | Energy cost of fat turnover |
| $\eta_{FFM}$ | 230 kcal/kg | Energy cost of protein turnover |
| $\rho_{FM}$ | 9300 kcal/kg | Energy density of fat mass |
| $\rho_{FFM}$ | 1100 kcal/kg | Energy density of fat free mass |
| $\beta$ | 0.24 | Dietary and adaptive thermogenesis |
| $UGE$ | 360 kcal/d | Energy loss due to increased urinary glucose excretion with canagliflozin (300 mg/d) |

In addition to validating our mathematical model using body composition and energy expenditure data from controlled feeding studies (20, 21, 22), the model has also demonstrated accurate weight loss predictions regardless of medication usage or comorbid conditions, including type 2 diabetes, in free-living individuals with obesity following reduced calorie diets (28).

### Modeling feedback control of energy intake

While many hormonal and neuronal factors are known to be involved in the regulation of food intake (3, 5), general properties of the feedback relationship between changes in body weight and changes in energy intake have not been well characterized or quantified. We tested the ability of two potential feedback models to describe the observed body weight profiles in response to sustained canagliflozin treatment.

In the first model, referred to as “proportional control”, changes in the signals regulating food intake depend only on the current body weight and are not dependent on the duration or rate of weight loss. In this model, the aggregate effect of different feedback signals regulating body weight are described by the equation:

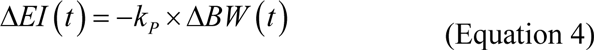

where the parameter *k*_*P*_ >0 quantifies the feedback strength. To simulate how proportional control of energy intake affects body weight kinetics during canagliflozin treatment, we added a UGE term to a validated mathematical model of adult human body weight dynamics (22) and calculated the mean Δ*EI* time course during placebo treatment to capture the typical transient weight loss effect of being in the trial (see (**Figure 1**). Equation 4 was then added to the placebo energy intake to simulate the mean homeostatic proportional control of Δ*EI* during canagliflozin treatment. The best fit parameter *k*_*P*_ was determined by a downhill simplex algorithm (29) implemented using Berkeley Madonna software (version 8.3; http://www.berkeleymadonna.com) to minimize the sum of squares residuals between the simulation outputs and the measured mean body weight and the calculated mean Δ*EI*.

**Figure 1.**
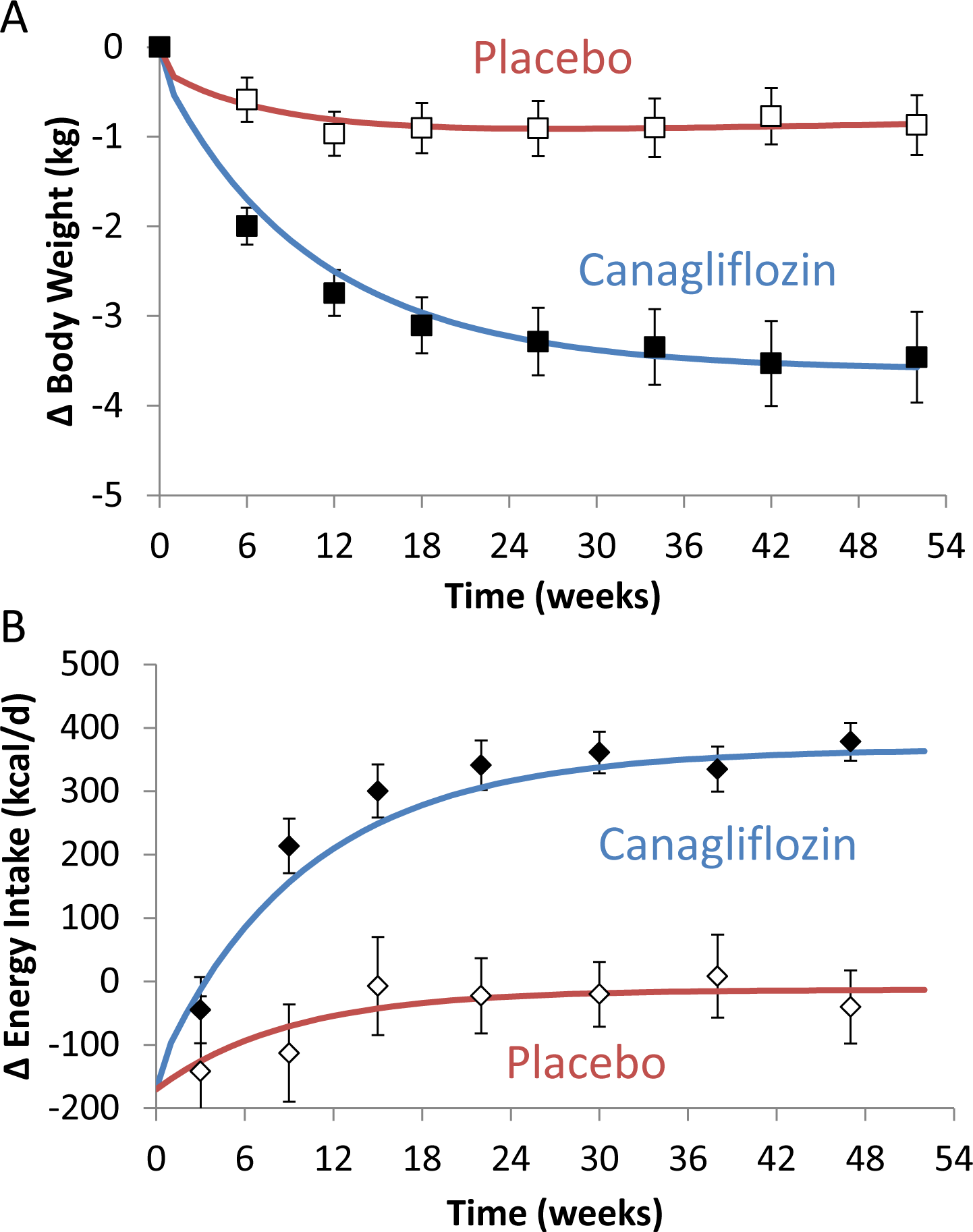
Body weight and energy intake changes during placebo and SGLT2 inhibition. (A) Average body weight measurements in the placebo group (□) and SGLT2 inhibition group (◼) along with mathematical model simulations depicted as red and blue curves, respectively. (B). Calculated energy intake changes in the placebo group (⋄) and the SGLT2 inhibitor group (♦) along with the mathematical model simulations. Mean ± 95% CI.

We also investigated another possible model for homeostatic body weight regulation that was previously suggested (30) such that changes in energy intake also depend on how long body weight has deviated from baseline, which is expressed as an integral term quantified by a parameter *k*_*I*_ >0:

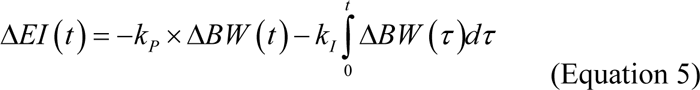

The integral feedback control was simulated using Equation 5 with the best fit value of *k*_*P*_ from Equation 4 and *k*_*I*_ = 1 kcal/kg/d^2^.

## Results

**Table 2** represents the characteristics of the type 2 diabetic subjects who completed the 52 week study and had body weight measurements throughout. The full study cohort was previously reported (18). In response to the sustained increase in UGE with canagliflozin treatment, mean body weight declined and reached a new equilibrium several kilograms lower and significantly more than the placebo group whose mean body weight loss was less than 1 kg ((**Figure 1A**). To explain the measured body weight changes in the treatment group given the estimated increases in UGE, energy intake was calculated to have increased by ~350 kcal/day at steady state ((**Figure 1B**) which is similar to recent estimates of the mean energy intake changes during 90 weeks of empagliflozin treatment, another SGLT2 inhibitor (31). In the placebo group, mean energy intake was calculated to transiently decrease by ~100 kcal/day over the first several weeks and return to baseline after 15 weeks (**Figure 1B**)

**Table 2.** Baseline characteristics of the study subjects. Data are mean ± SD unless otherwise indicated. ^†^ Includes American Indian or Alaska Native, Native Hawaiian or other Pacific Islander, multiple, other or not reported.

| Characteristic | Placebo<br>(n = 89) | Canagliflozin<br>(n = 153) |
| --- | --- | --- |
| Sex, n (%) |  |  |
| Male | 47 (53) | 66 (43) |
| Female | 42 (47) | 87 (57) |
| Age (years) | 57 ± 10 | 55 ± 11 |
| BW (kg) | 88 ± 18 | 87 ± 21 |
| BMI (kg/m<sup>2</sup>) | 32 ± 6 | 32 ± 6 |
| Waist circumference (cm) | 106 ± 13 | 105 ± 15 |
| Race, n (%) |  |  |
| White | 62 (70) | 106 (69) |
| Black or African American | 2 (2) | 7 (5) |
| Asian | 15 (17) | 24 (16) |
| Other<sup>†</sup> | 10 (11) | 16 (10) |
| HbA1c (%) | 7.5 ± 0.6 | 7.9 ± 0.8 |
| Fasting plasma glucose (mmol/L) | 8.2 ± 2.1 | 9.3 ± 2.4 |
| Diabetes duration (y) | 3 ± 4 | 3 ± 4 |
| eGFR (mL/min/1.73m<sup>2</sup>) | 85 ± 21 | 88 ± 19 |

Note that as body weight declined during SGLT2 inhibition energy intake increased until it compensated for the loss of calories via UGE, after which, a new equilibrium was reached. This general pattern of response, in which a sustained perturbation (caloric loss via UGE) leads to a new equilibrium at a lower body weight, suggests that the endocrine signals responding to weight loss (such as leptin) act as part of a proportional feedback control system as shown in **Figure 2** illustrating that proportional feedback control (as defined by Equation 4 with the parameter *k*_*P*_ = 95 kcal/day per kg) mimics the observed body weight and energy intake patterns in humans receiving SGLT2 inhibitors.

**Figure 2.**
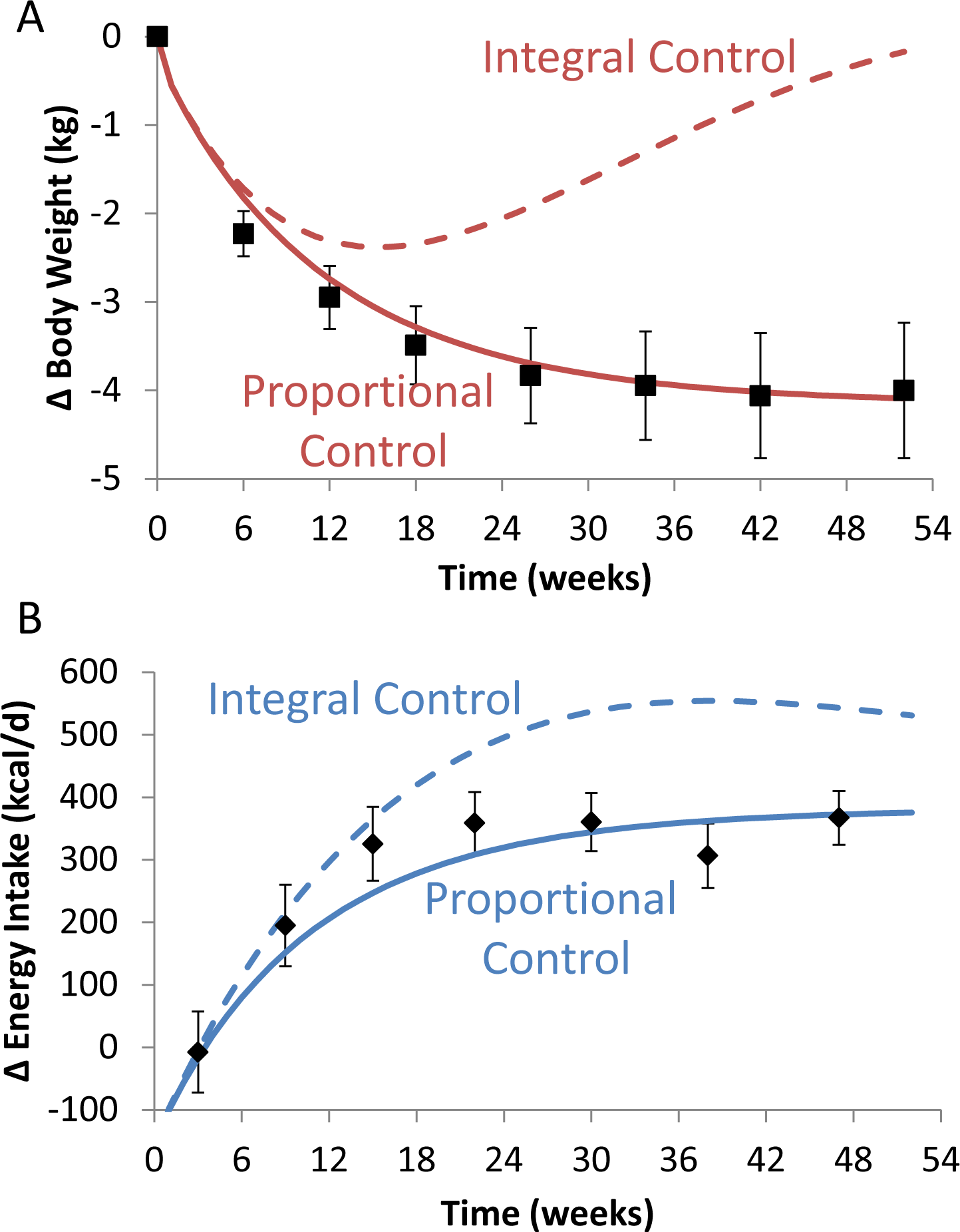
Characterization of feedback control of energy intake in subjects with type 2 diabetes treated with canagliflozin (18). (A) Observed changes in body weight (◼) and the simulated changes with proportional control (solid red curve) or integral control (dashed red curve) of energy intake. (B) Calculated changes in energy intake (♦) and the simulated changes with or proportional control (solid blue curve) integral control (dashed blue curve) of energy intake. Mean ± 95% CI.

In contrast, despite the persistent increase in UGE, if the system regulating body weight included integral feedback control (Equation 5), body weight would ultimately have been restored to baseline values even in the presence of sustained increases in UGE. This outcome is a well-known result from control theory that integral feedback is a necessary and sufficient condition for a system to produce zero steady-state error (i.e., return to baseline body weight) in response to a sustained perturbation (i.e., persistently increased UGE) and that a proportional feedback system will always have a non-zero steady-state error (i.e., a sustained reduction in body weight) in response to a sustained perturbation (32). In contrast, for any value of *k*_*I*_ > 0, a system including integral feedback in the regulation of energy intake would have only transient weight loss during sustained SGLT2 inhibitor treatment as shown for the integral feedback model in Figure 2.

By modeling the mean changes in energy intake obtained with SGLT2 inhibitor treatment using the proportional feedback model, our results quantify the strength of homeostatic energy intake control in humans. On average, energy intake increased by ~100 kcal/day per kg of weight lost—an effect substantially greater than the ~30 kcal/kg/day changes in energy expenditure observed with 10 to 20% weight loss in subjects with obesity (1).

To put our results into context, consider the body weight trajectory depicted in **Figure 3A** that resulted from long-term participation in a structured commercial weight loss program (33) and illustrates the ubiquitous body weight time course characterized by initial weight loss, a plateau after 6-8 months, followed by slow weight regain (34). The calculated energy intake corresponding to this mean body weight trajectory is illustrated in **Figure 3B** showing an initial reduction of ~700 kcal/day from baseline followed by an exponential relaxation towards baseline over the ensuing months. The estimated energy expenditure corresponding to this intervention is also shown in Figure 3B and indicates relatively minor changes in comparison to the energy intake changes. Note that at the point of maximum weight loss occurring at the ~8 month plateau, energy intake had already returned to within 100 kcal/day of baseline. After 1 year, the average energy intake was practically at baseline levels while body weight was still reduced by ~5 kg. A similar pattern of relapsing diet adherence has been observed using objective biomarker methods during an intensive 2 year calorie restriction study where calorie intake was much lower during the early period of weight loss than after the body weight had plateaued (16, 35).

**Figure 3.**
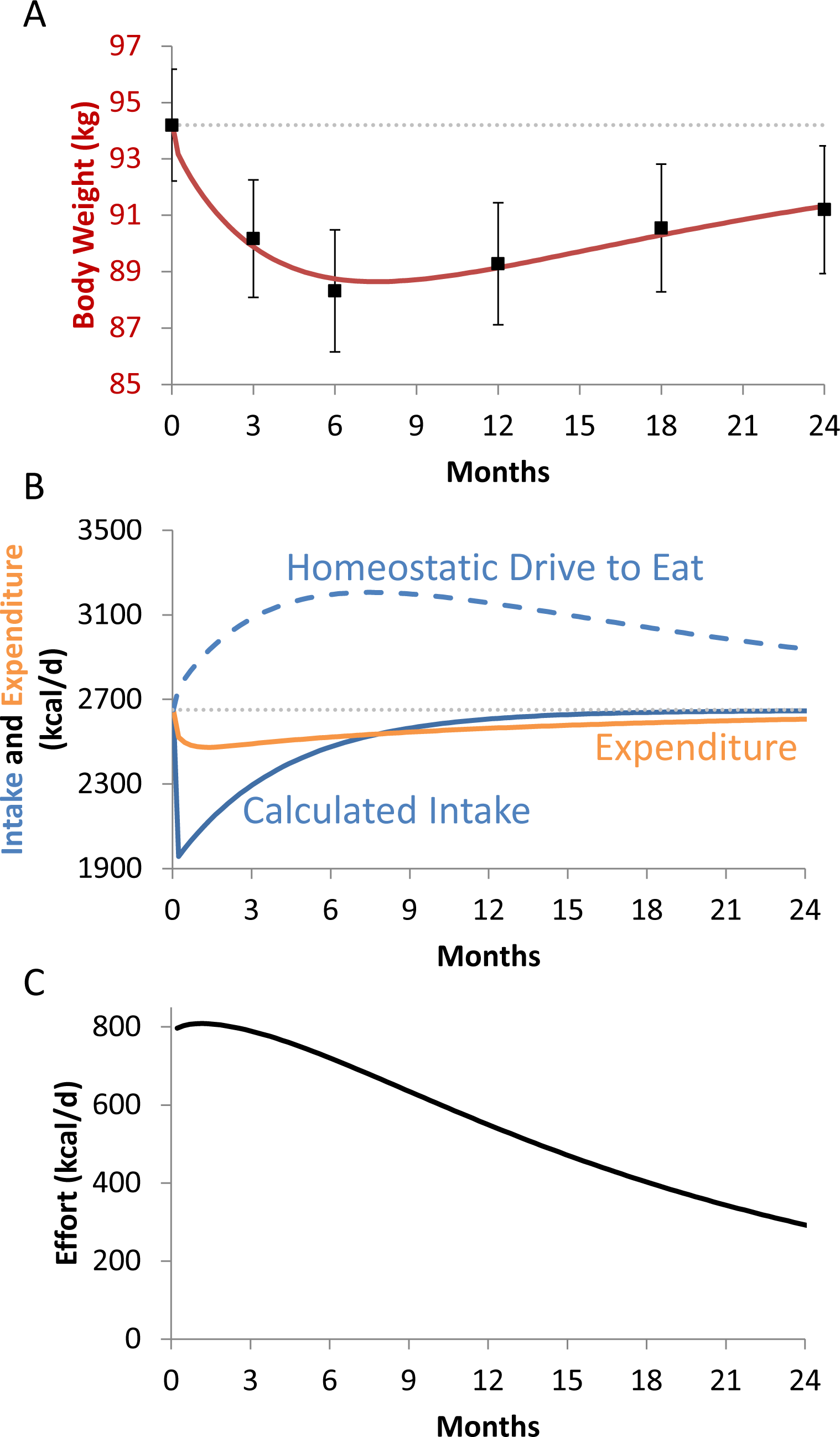
Energy balance dynamics during a lifestyle intervention for weight loss (33). (A) Average body weight (◼) typically decreases and reaches a plateau after 6–8 months of a lifestyle intervention followed by slow weight regain. (B) Energy expenditure changes relatively little during the intervention (solid orange curve) whereas energy intake initially drops by a large amount followed by an exponential return towards baseline (solid blue curve). The homeostatic feedback from the body weight loss signals a large increase in homeostatic drive to eat (dashed curve) that is resisted by the attempt to sustain the intervention. (C) The average effort during the intervention was defined as the difference between the homeostatic drive to eat and the actual energy intake. A
substantial effort persists during the intervention despite a return to near baseline energy intake. Mean ± 95% CI.

The calculated energy intake time course during the lifestyle intervention suggests that diet adherence lapsed very early with subjects returning to their previous caloric intake and a corresponding weight plateau followed by slow weight regain. While this might be interpreted as indicating that the participants rapidly decreased their effort to adhere to the intervention, it is enlightening to consider their food intake behavior within the context of the proportional feedback control system. The dashed curve in Figure 3B illustrates the energy intake pattern corresponding to the increased homeostatic drive to eat in response to the body weight changes shown in Figure 3A. The difference between the homeostatic drive to eat and the actual energy intake is depicted in Figure 3C and is a quantitative index of the ongoing effort to sustain the intervention in the face of the continuing biological signals to overeat. In this context, a substantial persistent effort is required to avoid overeating above baseline during the intervention despite the average energy intake returning to near baseline levels.

## Discussion

In the absence of ongoing efforts to restrain food intake following weight loss, homeostatic feedback control of energy intake will result in eating above baseline levels with an accompanying acceleration of weight regain. Such behavior has been previously observed in rodent models when a return to *ad libitum* feeding following diet restriction resulted in hyperphagia until the lost weight was regained (36). This phenomenon has been also observed in lean men following experimental semi-starvation (37) or short-term underfeeding (38, 39, 40). Hyperphagia in these studies was believed to result from homeostatic signals arising from loss of both body fat and lean tissues (41, 42), but a conscious desire to regain lost weight cannot be ruled out and may have contributed to the increased food intake.

Previous studies of energy intake regulation in humans have employed short-term diet manipulations to measure compensatory changes in energy intake (43, 44, 45, 46). While such studies can provide useful information about the influence of episodic appetite signals on short-term modulation of energy intake, the results cannot be readily extrapolated to the long time scales associated with regulation of human energy balance and do not provide information about how weight changes influence energy intake.

Long-term inhibition of SGLT2 provides a unique probe for assessment of human energy homeostasis since the mechanism of action is clear, its effect on energy output is consistent, and the intervention is unlikely to directly affect central pathways involved in regulation of food intake. In contrast, other interventions aimed at increasing energy expenditure, such as exercise (17) or exogenous delivery of thyroid hormone (47), have pleiotropic effects and their impact on energy expenditure can be highly variable.

The suggestion that the signals controlling energy intake act as a proportional feedback system without integral feedback is consistent with the roughly proportional changes in appetite-regulating hormones that occur rapidly in response to weight loss and do not further increase as weight loss is sustained (as would occur with integral feedback) (48). We do not yet know whether the simple proportional controller represented by Equation 4 is valid for a range of weight losses. For example, it may be possible that small weight changes are uncompensated by changes in energy intake such that the control system engages only after sufficient weight loss to cross some threshold (49). Furthermore, larger weight losses may result in energy intake adaptations corresponding to a nonlinear function of body weight change. Future research is required to address these questions and more fully characterize the homeostatic energy intake control system, characterize its variability between individuals, and identify its physiological mediators in humans.

Proportional homeostatic control of energy intake may help explain why the calculated exponential decay of diet adherence during weight loss interventions markedly contrasts with self-reported measurements that indicate persistence of diet adherence and no significant differences in caloric consumption between the period of early weight loss compared with the later time when weight has plateaued (50). This has led to speculation that the 6-8 month weight plateau may be entirely due to slowing of metabolic rate rather than loss of diet adherence (51). Our results suggest otherwise and self-reported energy intake measurements are well-known to be quantitatively unreliable (13). Nevertheless, the relative constancy self-reported energy intake over the first 6 months corresponds well with the calculated persistent effort to resist the homeostatic drive to overeat at above baseline levels. Therefore, self-reported measurements of diet may more accurately reflect the perceived effort of the dieter to adhere to the intervention rather than their actual energy intake.

An important limitation of our study is that all of the subjects had type 2 diabetes and it is unclear whether our results translate to people without diabetes. For example, people with type 2 diabetes have a characteristically decreased insulin secretion in the context of insulin resistance that may quantitatively affect the feedback control of energy intake (3, 4, 5). Another limitation is that we cannot be certain that weight loss achieved by mechanisms other than SGLT2 inhibition will lead to the same homeostatic response to increase energy intake. Furthermore, we restricted our analysis to the group mean rather than attempt to characterize individual responses since UGE was not directly measured in each subject but was assumed to be ~90 g/day based on previous measurements in people with type 2 diabetes treated with 300 mg/day canagliflozin (19, 52, 53). A subset of subjects in this study underwent a meal tolerance test where UGE was measured; their mean increase in UGE over the 3h meal test was 14.7 g (54); this is similar to the value observed following meals in other studies where mean 24h UGE was measured to be ~90 g/day and suggests that the mean daily UGE in this study is unlikely to be substantially different than what was assumed. Therefore, there is a correspondingly small uncertainty in the mean calculated proportional control of energy intake (~100 kcal/day per kg of body weight lost) at the group level. Measuring daily UGE in individuals during long-term studies with SGLT2 inhibitors would enable the individual subject variability in the magnitude of the compensatory energy intake responses to be characterized.

In summary, our results provide the first quantification of homeostatic feedback control of energy intake in free-living humans. We found that appetite proportionately increased by ~100 kcal/day above baseline for every kilogram of lost weight. In comparison, energy expenditure adaptations to weight loss are several-fold lower in magnitude. Therefore, homeostatic feedback control of energy intake is likely a primary reason why it is so difficult to achieve large sustained weight losses in patients with obesity. Rather, weight regain is typical in the absence of heroic and vigilant efforts to maintain behavior changes in the face of an omnipresent obesogenic environment (55). Unfortunately, there is no evidence that the energy intake feedback control system resets or relaxes with prolonged maintenance of lost weight – an effect similar to the long-term persistent suppression of energy expenditure in weight-reduced humans (56). Therefore, the effort associated with a weight loss intervention persists until either body weight is fully regained or energy intake increases above baseline to match the homeostatic drive to eat. Permanently subverting or countering this feedback control system poses a major challenge for the development of effective obesity therapies.

## Acknowledgements

Canagliflozin has been developed by Janssen Research & Development, LLC, in collaboration with Mitsubishi Tanabe Pharma Corporation. We thank Marc L. Reitman and Aaron Cypess for their helpful comments on the manuscript.

DP and KDH had full access to all the data in the study and take responsibility for the integrity of the data and the accuracy of the data analysis. DP, KDH and RS designed the study; AS, DP, and KDH performed the analyses; DP, KDH, and RS interpreted the data and drafted the manuscript.

These data were previously presented, in part, in abstract form at the International Congress on Obesity on May 2, 2016 in Vancouver, Canada and at the 50th Annual Meeting of the European Association for the Study of Diabetes on September 15-19, 2014 in Vienna, Austria.

